# The environmental stress response causes ribosome loss in aneuploid yeast cells

**DOI:** 10.1101/2020.03.25.008540

**Authors:** Allegra Terhorst, Arzu Sandikci, Abigail Keller, Charles A. Whittaker, Maitreya J. Dunham, Angelika Amon

## Abstract

Aneuploidy, a condition characterized by whole chromosome gains and losses, is often associated with significant cellular stress and decreased fitness. However, how cells respond to the aneuploid state has remained controversial. In aneuploid budding yeast, two opposing gene expression patterns have been reported: the “environmental stress response” (ESR) and the “common aneuploidy gene-expression” (CAGE) signature, in which many ESR genes are oppositely regulated. Here, we investigate and bring clarity to this controversy. We show that the CAGE signature is not an aneuploidy-specific gene expression signature but the result of normalizing the gene expression profile of actively proliferating aneuploid cells to that of euploid cells grown into stationary phase. Because growth into stationary phase is amongst the strongest inducers of the ESR, the ESR in aneuploid cells was masked when stationary phase euploid cells were used for normalization in transcriptomic studies. When exponentially growing euploid cells are used in gene expression comparisons with aneuploid cells, the CAGE signature is no longer evident in aneuploid cells. Instead, aneuploid cells exhibit the ESR. We further show that the ESR causes selective ribosome loss in aneuploid cells, providing an explanation for the decreased cellular density of aneuploid cells. We conclude that aneuploid budding yeast cells mount the ESR, rather than the CAGE signature, in response to aneuploidy-induced cellular stresses, resulting in selective ribosome loss. We propose that the ESR serves two purposes in aneuploid cells: protecting cells from aneuploidy-induced cellular stresses and preventing excessive cellular enlargement during slowed cell cycles by downregulating translation capacity.

## Introduction

Dividing cells rely on multiple complex mechanisms to correctly segregate their chromosomes and create euploid progeny. When chromosome missegregation occurs, daughter cells can acquire an incorrect number of chromosomes that is not a complete multiple of the haploid genome, a condition termed aneuploidy. Aneuploidy can occur naturally; for example, 17% of wild budding yeast isolates harbor aneuploidies and are thought to have evolved mechanisms to tolerate these aneuploid karyotypes (1, 2). In most cases, however, aneuploidy is highly detrimental, especially in multicellular animals (3).

Various models have been developed to study aneuploidy in *S. cerevisiae*. Their analyses led to the conclusions that aneuploidy affects a wide range of cellular processes, such as protein homeostasis, metabolism, and cell wall integrity, and results in an overall decrease in cellular fitness (3, 4). However, how aneuploidy affects gene expression has remained controversial. While it is clear that gene expression scales with gene copy number in aneuploid cells, there is not yet a consensus on whether aneuploidy elicits a global transcriptional response in yeast and what this response may be.

We previously described that haploid aneuploid yeast cells harboring only one additional chromosome (henceforth disomic yeast strains) experience an environmental stress response (ESR) (5). The ESR is a transcriptional signature observed in response to nearly every type of exogenous stress, including hyper-osmotic, heat shock, oxidative and reductive stress, and nutrient limitation. These conditions cause the coordinated upregulation of approximately 300 genes, also known as the “induced (i)ESR” and downregulation of approximately 600 genes, also known as the “repressed (r)ESR” (6, 7). Genes that are upregulated compensate for various stressors and encode chaperones, amino acid transporters, and proteins involved in increasing endocytosis and proteasome activity. Downregulated genes encode factors critical for transcription and translation, among them are genes encoding ribosomal proteins and proteins involved in ribosome biogenesis (5, 6). The ESR is not only observed in response to stress but also in cells that grow slowly or cells that are cell cycle-arrested (4, 5, 8). Indeed, the strength of the ESR, that is the degree to which iESR genes are upregulated and rESR genes are downregulated, correlates remarkably well with growth rate, suggesting that this transcriptional signature is primarily determined by proliferation rate (4).

A recent study by Tsai et al. (2019; 9) reported that yeast cell populations harboring heterogenous aneuploidies do not exhibit the ESR. Instead, these aneuploid populations were described to exhibit a transcriptional response, termed the “common aneuploidy gene-expression” (CAGE) response. In the CAGE response, the genes that are upregulated in the ESR are downregulated, and those that are downregulated in the ESR are upregulated. The authors further found that the CAGE response bears similarity to a hypo-osmotic shock gene expression pattern, which was proposed to counter a decrease in cytoplasmic density observed in aneuploid cells (9, 10).

We report here the reanalysis of the gene expression data generated by Tsai et al. (2019; 9) as well as replication of their experimental approach. These analyses showed that the CAGE gene expression signature described by Tsai et al. (2019; 9) is an artifact caused by normalizing the gene expression of actively dividing aneuploid cells to that of euploid control cells that had grown to stationary phase. Growth into stationary phase is amongst the strongest inducers of the ESR (6). Thus, when Tsai et al. (2019; 9) compared the gene expression pattern of euploid stationary phase cells to that of aneuploid cells that, due to their poor proliferation, had not yet reached stationary phase, the ESR caused by aneuploidy was obscured. We find that when exponentially growing euploid cells were used in gene expression comparisons with aneuploid cells, the CAGE signature of aneuploid cells is no longer evident. Instead, aneuploid cell populations are found to exhibit the ESR, confirming previous reports (5). Using strains harboring multiple aneuploidies, we further show that the ESR causes a selective loss of ribosomes in aneuploid cells, providing a potential explanation for decreased cellular density previously reported to occur in response to chromosome gains and losses. We conclude that aneuploid budding yeast cells mount the ESR in response to aneuploidy-induced cellular stresses that results in ribosome loss.

## Results

### Exponentially growing haploid cells exhibit a transcriptional response, previously described to be unique to aneuploid cells

A recent study (9) reported the absence of the environmental stress response (ESR) in populations of yeast cells harboring different, random aneuploid karyotypes. Instead, it was reported that these heterogeneous aneuploid yeast populations exhibit the “common aneuploidy gene-expression” (CAGE) signature. In this study, Tsai et al. (2019; 9) developed two protocols to generate heterogeneous populations of aneuploid cells, taking advantage of the fact that sporulation of triploid cells results in high levels of aneuploid progeny. In the first method, Tsai et al. (2019; 9) dissected spores obtained from triploid cells, grew individual aneuploid spores into colonies, pooled these colonies and analyzed the gene expression pattern of these cells (*SI Appendix*, Fig. S1*A*). We will refer to these aneuploid populations as “aneuploid populations obtained from tetrads”. Euploid haploid cells obtained from sporulating diploid cells and handled in the same manner as aneuploid cells served as the control (henceforth “euploid populations obtained from tetrads”). In the second protocol, Tsai et al. (2019; 9) sporulated triploid cells, and then selected viable *MATa* aneuploid colonies by selecting for histidine prototrophy brought about by *HIS5* expressed from the *MATa*-specific *STE2* promoter (*SI Appendix*, Fig. S1*A*). We will refer to these aneuploid populations as “aneuploid populations obtained from *MATa* selection”. Again, euploid haploid cells obtained by sporulating diploid cells and *MATa* selected served as the euploid control (henceforth “euploid populations obtained from *MATa* selection”). Gene expression analysis of these cell populations led to the identification of an expression signature Tsai et al. (2019; 9) termed “common aneuploidy gene-expression” (CAGE) response. This gene expression signature resembles a hypo-osmotic stress response and is essentially oppositely regulated to the ESR; 59.8% of genes upregulated in the CAGE response are downregulated in the ESR, while 13.2% of CAGE downregulated genes are upregulated in the ESR (9).

Having previously identified the ESR in yeast strains harboring defined aneuploidies (4, 5), we wished to determine why pooled aneuploid populations did not exhibit the ESR but instead the CAGE gene expression signature. To this end, we reanalyzed the gene expression data reported by Tsai et al. (2019; 9) by individually processing the samples rather than normalizing the aneuploid cell populations to the euploid control populations. Among the RNA-Seq data sets deposited by Tsai et al. (2019; 9) was one obtained from an exponential growing haploid strain (RLY4388, accession number: GSM2886452 and GSM2886453), which the authors did not analyze. Using the RNA-Seq by Expectation Maximization (RSEM) processing method, we calculated the raw transcripts per million (TPM) values for the aneuploid and euploid cell populations as well as the exponentially growing haploid strain RLY4388 separately, then log_2_ transformed these values with a +1 offset to avoid negative expression values, and created row-centered heatmaps for genes upregulated and downregulated in both the CAGE and ESR gene expression signature (Fig. 1*A*). As previously reported, we observed the CAGE gene expression signature in the pooled aneuploid populations. Surprisingly, the exponentially growing haploid strain RLY4388, which was not analyzed by Tsai et al. (2019; 9), exhibited the strongest CAGE gene expression signature (Fig. 1*A*). Equally surprising was the observation that pooled euploid populations, used as normalization controls by Tsai et al. (2019; 9), exhibited the strongest ESR gene expression signature (Fig. 1*A*).

**Figure 1.**
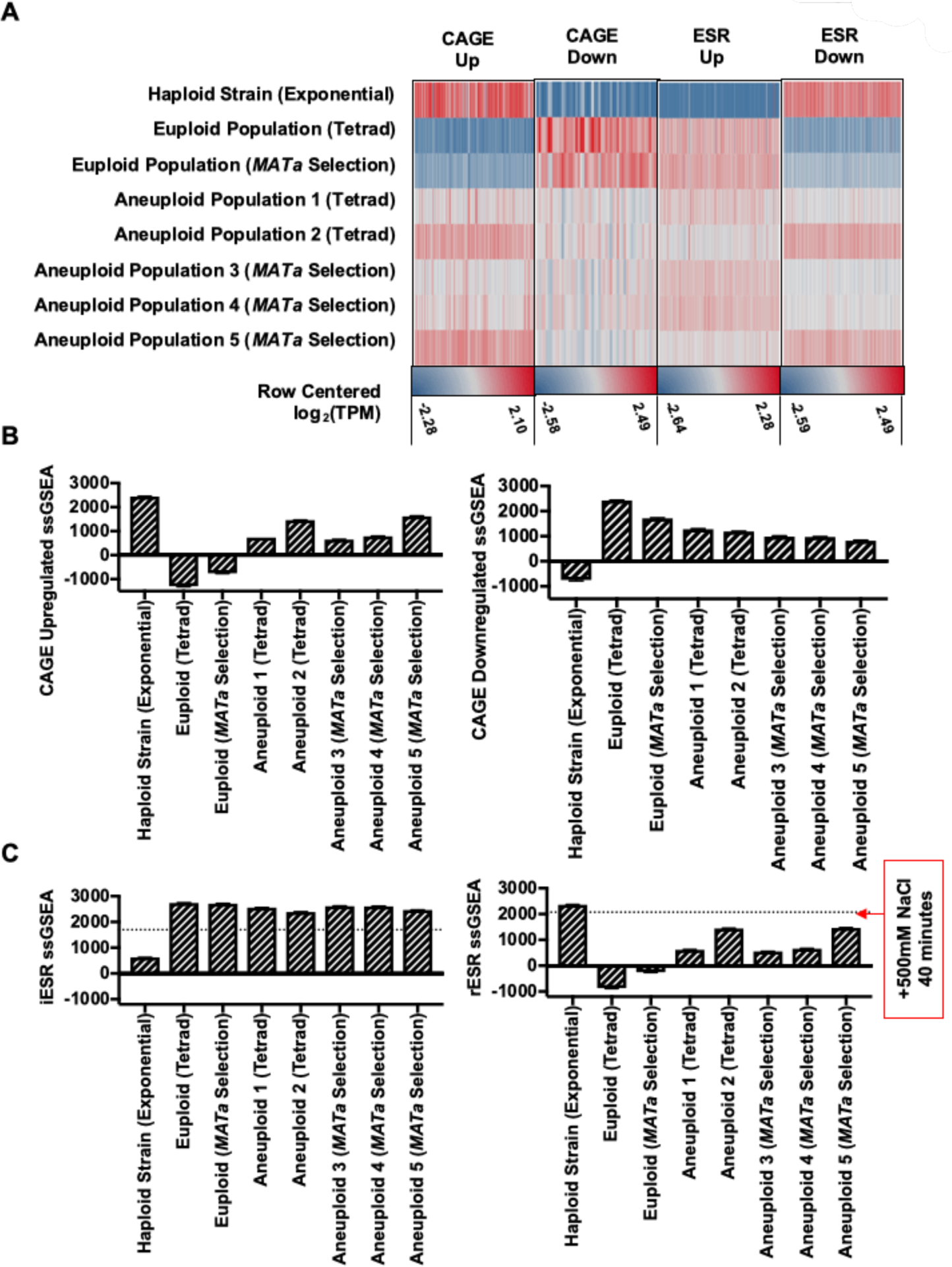
Reanalysis of published aneuploid transcription data from Tsai et al. (2019; 9). Transcription data of an exponentially growing haploid strain RLY4388 and euploid and aneuploid cell populations obtained from tetrad dissection (Tetrad) or *MATa* selection (*MATa* Selection) were reanalyzed with the RNA-Seq by Expectation Maximization (RSEM) processing method (Tsai et al. (2019; 9), accession number: GSE107997). Raw transcript per million (TPM) values were calculated for euploid cell populations, aneuploid cell populations, and the exponentially growing haploid strain. **(*A*)** Row centered log_2_(TPM) values for each gene expression set (CAGE upregulated, CAGE downregulated, iESR, and rESR). Each gene set was row centered individually and has a separate maximum (red) and minimum (blue) values, noted underneath. **(*B*)** CAGE upregulated and downregulated ssGSEA projection values for the exponentially growing haploid strain and euploid and aneuploid cell populations (Tetrad and *MATa* Selection). **(*C*)** iESR and rESR ssGSEA projection values for the exponentially growing haploid strain and euploid and aneuploid cell populations (Tetrad and *MATa* Selection). The horizontal lines represent the iESR and rESR ssGSEA projection values for W303 wild-type cells (A2587) treated with 500 mM NaCl for 40 minutes, a positive control for the ESR induction. Error bars represent standard deviation from the mean.

Tsai et al. (2019; 9) discovered the CAGE response and the absence of the ESR in aneuploid cell populations by normalizing the gene expression of aneuploid cell populations to euploid control cell populations (9). Given that our analysis of their raw data showed that the euploid control populations strongly exhibited the ESR, we used the gene expression data set obtained from the exponentially growing haploid strain RLY4388 to normalize the gene expression data from aneuploid populations instead of normalizing the data to that of euploid control populations. When compared to the gene expression data generated from the exponentially growing haploid strain RLY4388, aneuploid populations obtained from tetrad dissection and *MATa* selection exhibited the ESR, and the CAGE signature was no longer evident (*SI Appendix*, Fig. S2*A* and *B*). When we measured the differential gene expression between euploid populations and the exponentially growing haploid strain RLY4388, it was also apparent that the euploid populations (Tetrad and *MATa* Selection) were experiencing the ESR (*SI Appendix*, Fig. S2*C* and *D*).

Given that the choice of euploid control (euploid populations versus an exponentially growing haploid wild-type strain RLY4388) made such a large difference in the experimental outcome, we decided to employ a data analysis method that does not depend on normalization. Single-sample gene set enrichment analysis (ssGSEA) generates a single projection value for a set of genes within a sample. These values can then be compared between samples in order to measure how the gene expression distribution of that gene set changes across an experiment (11). Using this approach, we confirmed that aneuploid cell populations exhibited the CAGE signature (Fig. 1*B*; Mean CAGE Upregulated ssGSEA = 1046 ± 419.5, Mean CAGE Downregulated ssGSEA = 1048 ± 169.7), while the euploid control cell populations did not (Fig. 1*B*; Mean CAGE Upregulated ssGSEA = -1031 ± 299.7, Mean CAGE Downregulated = 2069 ± 392). However, unexpectedly, the sample with the strongest CAGE signature, thought to be a characteristic of aneuploidy, was obtained from the exponentially growing haploid strain RLY4388 (Fig. 1*B*; Mean CAGE Upregulated ssGSEA = 2438 ± 12.37, Mean CAGE Downregulated ssGSEA = -746.5 ± 32.61).

ssGSEA analysis of the ESR in aneuploid and euploid cell populations revealed equally unanticipated results. As expected, the exponentially growing haploid strain RLY4388 did not exhibit the ESR (Fig. 1*C*; Mean iESR ssGSEA = 621.9 ± 3.596, Mean rESR ssGSEA = 2362 ± 5.558). Consistent with our previous observations in disomic yeast strains (5), aneuploid cell populations showed the ESR (Fig. 1*C*; Mean iESR ssGSEA = 2525 ± 89.96, Mean rESR ssGSEA = 947.7 ± 428.4), but the euploid control populations exhibited an even stronger ESR (Fig. 1*C*; Mean iESR ssGSEA = 2723 ± 15.57, Mean rESR ssGSEA = -557.9 ± 338.9). The degree to which the ESR was induced in these euploid control populations was greater than in exponentially growing wild-type cells (A2587) treated with 500 mM NaCl for 40 minutes. We conclude that the euploid control populations analyzed by Tsai et al. (2019; 9) exhibit the strongest ESR signature, indicating that they experienced significant exogenous stress.

### Stationary phase cells exhibit the environmental stress response

It was curious that the euploid control populations generated by Tsai et al. (2019; 9) strongly exhibited the ESR. To determine the cause of this robust expression of the ESR, we repeated their tetrad dissection protocol to obtain euploid and aneuploid cell populations, employing the strains used by Tsai et al. (2019; 9) after detailed consultation with the authors. We dissected 200 and 770 tetrads obtained from diploid and triploid cells, respectively. Spore viability for the euploid strain was 97.3% and, as expected, significantly lower for triploid strains (40.2%) because many aneuploid strains are inviable. We then followed the protocol developed by Tsai et al. (2019; 9) and grew colonies obtained from viable spores in individual wells of a 96-well deep well plate for 14-16 hours in 200 μL YEPD medium at 25 °C (*SI Appendix*, Fig. S1*A*) Thereafter, we added 300 μL YEPD medium to cultures and grew them for an additional 5 hours at 25 °C. The euploid and aneuploid cultures were then separately pooled to create heterogeneous euploid and aneuploid cell populations. Using this growth protocol, pooled euploid populations had reached an OD(600nm) of 8.54. As expected, owing to aneuploid cells having significant proliferation defects, pooled aneuploid populations reached an OD(600nm) of only 2.62.

The high OD(600nm) values reached by the euploid population provided a potential explanation for why they strongly exhibited the ESR. As cultures approach stationary phase, cells experience starvation, which is among the strongest inducers of the ESR (6). To test this hypothesis, we determined at which OD(600nm) S288C wild-type haploid cells activate the ESR. We grew cells into stationary phase in YEPD medium and measured rESR and iESR gene expression over time (Fig. 2*A*). iESR gene induction was observed at around an OD(600nm) of 3.5, determined by an increase in iESR ssGSEA projection values; rESR gene expression began to decline dramatically at an OD(600nm) of 5.5. These results explain why euploid control populations analyzed by Tsai et al. (2019; 9) exhibit such a strong ESR but aneuploid populations did not. The aneuploid cells had not yet reached an OD(600nm) value where starvation-induced ESR induction occurs.

**Figure 2.**
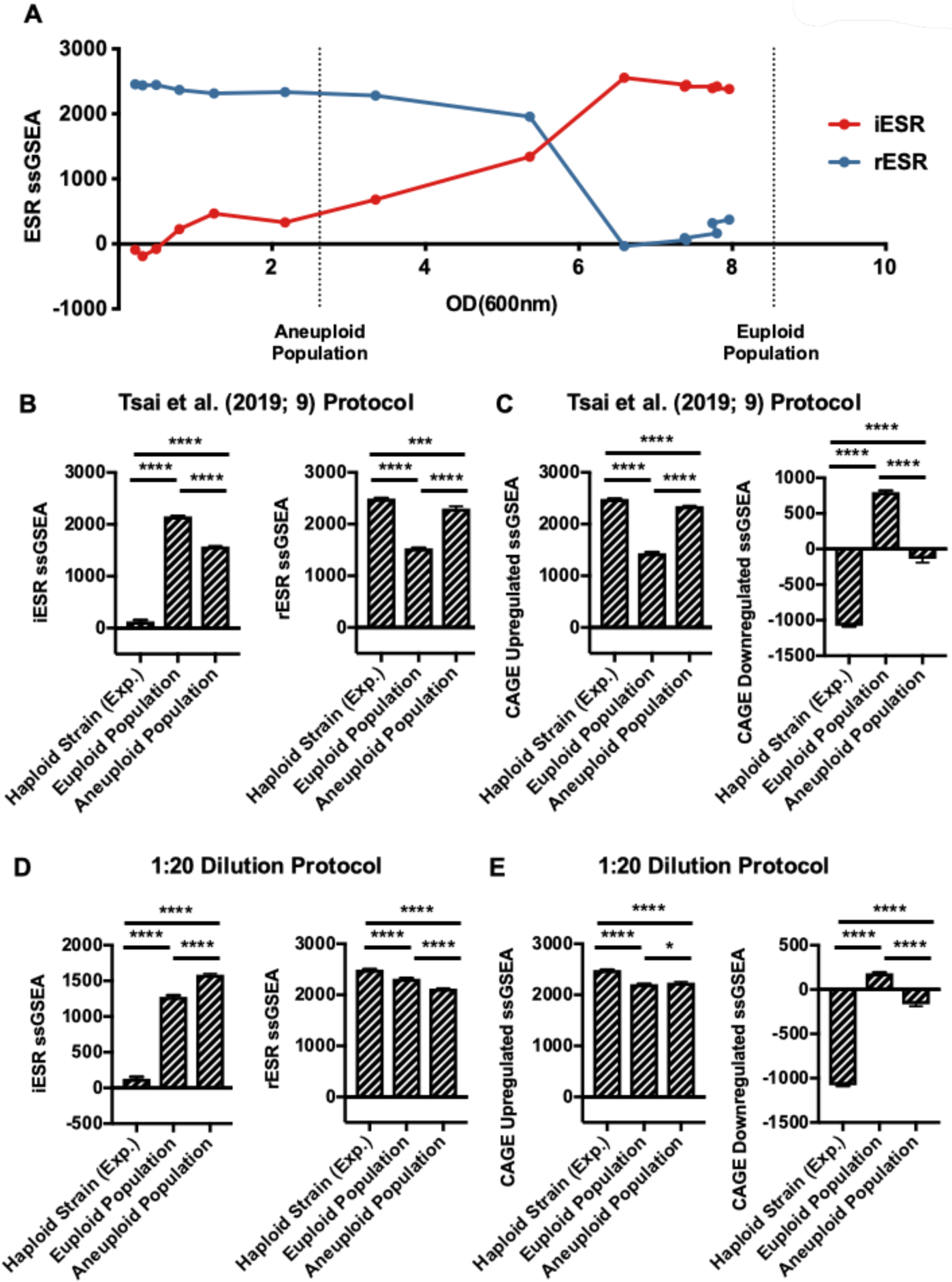
Effects of cell density on ESR strength in aneuploid cell populations. **(*A*)** iESR (red) and rESR (blue) ssGSEA projection values were determined at the indicated OD(600nm) for S288C wild-type haploid cells (A2050) grown in YEPD over 28 hours. Vertical lines represent the OD(600nm) values of pooled euploid and aneuploid cell populations generated by tetrad dissection. Error bars represent standard deviation from the mean. **(*B* and *C*)** Tetrads of sporulated S288C diploid and triploid cells (A40877, A40878) were dissected to produce heterogeneous haploid and aneuploid cell populations, respectively. 144 individual haploid colonies and 432 aneuploid colonies were inoculated and grown overnight in 200 μL YEPD. The next morning 300 μL YEPD were added to cultures and grown for an additional 5 hours. Individual euploid and aneuploid cultures were then pooled and their transcriptomes analyzed. An exponentially growing haploid strain (A2050) was included as a control. Gene expression data were analyzed by calculating ssGSEA projection values for the **(*B*)** iESR and rESR and **(*C*)** CAGE upregulated and downregulated genes. Error bars represent standard deviation from the mean; one-way ANOVA test with multiple comparisons and Bonferroni correction, *P* < 0.0001 (****), *P* = 0.0002 (***). **(*D* and *E*)** Tetrads of sporulated S288C diploid and triploid cells (A40877, A40878) were dissected to produce heterogeneous haploid and aneuploid cell populations, respectively. 144 individual haploid colonies and 432 aneuploid colonies were inoculated and grown overnight in 200 μL YEPD. The next morning cultures were diluted 1:20 and grown for an additional 5 hours. Colonies were then pooled, further diluted to approximately OD(600nm) = 0.3, and grown for 2 additional hours. Transcriptomes of pooled euploid and aneuploid populations and an exponentially growing haploid strain (A2050) were analyzed with RNA-Seq, and ssGSEA projection values were calculated for **(*D*)** iESR and rESR and **(*E*)** CAGE upregulated and downregulated genes. Error bars represent standard deviation from the mean; one-way ANOVA test with multiple comparisons and Bonferroni correction, *P* < 0.0001 (****), *P* = 0.0332 (*).

To confirm this conclusion, we analyzed the gene expression profile in our euploid and aneuploid cell populations grown using the protocol employed by Tsai et al. (2019; 9). Euploid control populations exhibited a stronger ESR than aneuploid cell populations. The iESR ssGSEA for euploid strains was 2155 ± 4.394 compared to an iESR ssGSEA of 1476 ± 11.18 in aneuploid populations. The rESR ssGSEA was 1533 ± 7.324 in euploids but 2300 ± 45.81 in aneuploids (Fig. 2*B*). Consistent with the idea that the CAGE signature is essentially the opposite of the ESR, aneuploid strains exhibited a stronger CAGE signature than euploid strains (Fig. 2*C*; aneuploid populations: CAGE Upregulated ssGSEA = 2348 ± 9.116, CAGE Downregulated ssGSEA = -136.6 ± 53.29; euploid populations: CAGE Upregulated ssGSEA = 1437 ± 19.08, CAGE Downregulated ssGSEA = 800.3 ± 19.29).

Our experiment also included a sample of an exponentially growing haploid wild-type strain (A2050). As expected, this strain did not exhibit the ESR (Fig. 2*B*) but instead showed the strongest CAGE response among all the cultures analyzed (Fig. 2*C*). Together, these data indicate that the CAGE response is not an aneuploidy-specific gene expression signature but the result of differences in proliferation rates between aneuploid and euploid cell populations. In the growth protocol employed by Tsai et al. (2019; 9), euploid cells had reached stationary phase, which causes a very strong ESR. In contrast, aneuploid cells had not. Because the ESR of aneuploid cells is weaker than that of stationary phase euploid cells, normalization of the aneuploid gene expression profile to that of stationary euploid cells led to the incorrect conclusion that aneuploid cells exhibited a transcriptional signature opposite of the ESR. This conclusion predicts that when growth into stationary phase is avoided, aneuploid cell populations ought to exhibit the ESR stronger than euploid control populations.

### Aneuploid cell populations exhibit the ESR

To determine whether growth of the control euploid population into stationary phase precluded the identification of the ESR in aneuploid cell populations, we repeated the protocol developed by Tsai et al. (2019; 9) to generate euploid and aneuploid cell populations. However, instead of diluting cultures 1:2 fold after 14–16 hours of growth in 200 uL of YEPD, we diluted cultures 1:20 fold (*SI Appendix*, Fig. S1*B*; henceforth 1:20 dilution protocol). This prevented either culture from reaching stationary phase, and the final OD(600nm) of pooled euploid and aneuploid populations was 0.29 and 0.3, respectively.

Gene expression analysis of these cultures resulted in a strikingly different outcome compared to that obtained from cells where euploid control populations had reached stationary phase. Aneuploid populations exhibited a stronger ESR than euploid control populations (Fig. 2*D*; iESR euploid population ssGSEA = 1277 ± 20.68, rESR ssGSEA = 2315 ± 11.89 compared to iESR ssGSEA = 1586 ± 9.762, rESR = 2122 ± 14.34 in aneuploid populations; iESR *P* < 0.0001, rESR *P* < 0.0001). It is, however, noteworthy that euploid control populations also exhibited the ESR, although not as strong as aneuploid populations, when compared to an exponentially growing wild-type strain (Fig. 2*D*; iESR ssGSEA = 129.6 ± 25.11, rESR ssGSEA = 2496 ± 7.945). This is likely due to the fact that euploid cell populations grown in deep wells experience nutrient limitation. Aeration is poorer and proliferation rate slower in deep well plates compared to in a vigorously shaking flask.

Analysis of the CAGE signature revealed that the exponentially growing wild-type strain expressed CAGE genes much more strongly than either the euploid (CAGE Upregulated *P* < 0.0001, CAGE Downregulated *P* < 0.0001) or aneuploid cell populations (CAGE Upregulated *P* < 0.0001, CAGE Downregulated *P* < 0.0001) (Fig. 2*E*). We note that the aneuploid population showed a slightly greater decrease in expression of the down-regulated CAGE response than euploid control populations (Fig 2*E*; aneuploid CAGE Downregulated ssGSEA = -166.0 ± 23.11 versus euploid CAGE Downregulated ssGSEA = 184.3 ± 8.571). While this difference is statistically significant (*P* < 0.0001), it is likely biologically irrelevant given the dramatically higher downregulation of CAGE genes in the exponentially growing haploid wild-type strain. We conclude that aneuploid cell populations exhibit the ESR and that the previously reported aneuploidy specific CAGE signature is most prominent in an exponentially growing haploid strain.

### Degree of aneuploidy correlates to ESR strength in complex aneuploid strains

Previous results from our lab indicated that yeast strains harboring an additional chromosome (disomes) activate the ESR, and our results shown here demonstrate that heterogeneous aneuploid populations do too (5). We next wished to determine whether this gene expression signature is also present in yeast strains harboring multiple specific aneuploidies. Pavelka et al. (2010; 12) created a large number of yeast strains carrying multiple aneuploidies by sporulating a pentaploid strain (12). Strains obtained from such spores harbor multiple aneuploidies ranging in genome content between 2N and 3N (*SI Appendix*, Table S2; 12). Because the strength of the ESR is largely defined by proliferation rate (4, 8), we first measured doubling times of these complex aneuploid strains to ask whether proliferation rate was correlated with degree of aneuploidy also in strains harboring multiple aneuploidies. We calculated degree of aneuploidy as the fraction of base pairs in the aneuploid strain vis-à-vis a haploid euploid control strain. We found that the proliferation defect of aneuploid strains correlated remarkably well with their degree of aneuploidy (Fig. 3*A*; Spearman, ρ^2^ = 0.7620, *P* < 0.0001).

**Figure 3.**
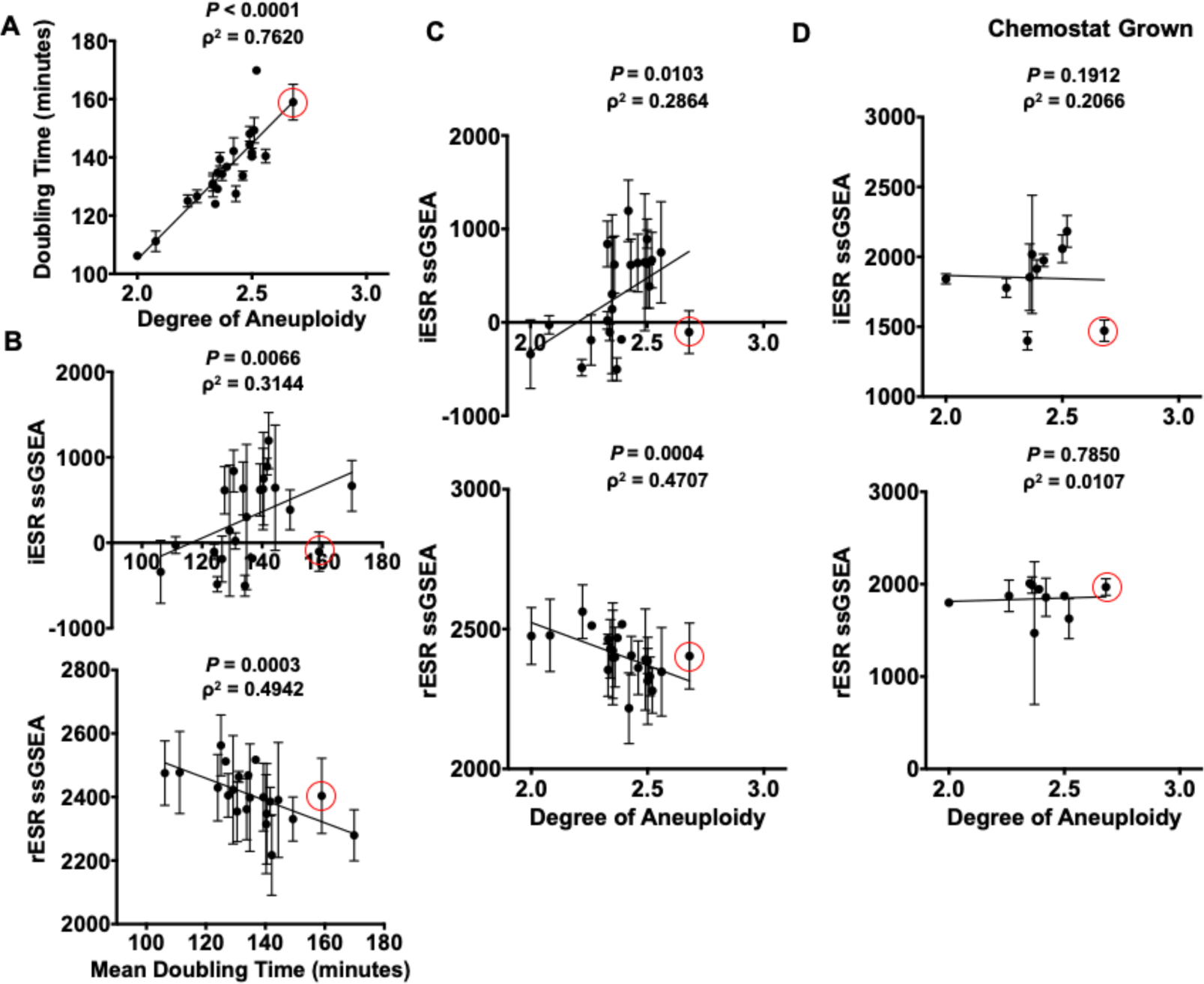
Complex aneuploid yeast strains exhibit the ESR. **(*A-C*)** Aneuploid yeast strains harboring aneuploidies ranging from 2N to 3N were grown to log phase in YEPD. For each strain, degree of aneuploidy was calculated as the fraction of base pairs in the aneuploid strain/base pairs in a haploid control strain. Doubling times were calculated from growth curves generated by measuring OD(600nm) in 20 minute intervals over 5 hours in a plate reader. **(*A*)** Correlation between doubling time and degree of aneuploidy (Spearman, ρ^2^ = 0.7620, *P* < 0.0001). Transcriptomes of the complex aneuploid strains were analyzed by RNA-Seq, and ssGSEA projection values were calculated for iESR and rESR genes. Correlations between iESR ssGSEA projections and mean doubling time (Spearman, ρ^2^ = 0.3144, *P* = 0.0066) and rESR ssGSEA projections and mean doubling time (Spearman, ρ^2^ = 0.4942, *P* = 0.0003) are shown in **(*B*)**. Correlations between iESR ssGSEA projections and degree of aneuploidy (Spearman, ρ^2^ = 0.2864, *P* = 0.0103) and rESR ssGSEA projections and degree of aneuploidy (Spearman, ρ^2^ = 0.4707, *P* = 0.0004) are shown in **(*C*)**. Error bars represent standard deviation from the mean. **(*D*)** Select complex aneuploid strains were grown in a phosphate-limiting chemostat until steady state was reached. Transcriptomes of harvested cells were analyzed by RNA-Seq. ssGSEA projection values were calculated for iESR and rESR genes. Correlations between iESR ssGSEA projections and degree of aneuploidy (Spearman, ρ^2^ = 0.1912, *P* = 0.2066) and rESR ssGSEA projections and degree of aneuploidy (Spearman, ρ^2^ = 0.0107, *P* = 0.7850) are shown. Error bars represent standard deviation from the mean. The data point circled in red represents a complex aneuploid strain that does not mount the ESR.

Gene expression analysis of these complex aneuploid strains further revealed a strong correlation between mean doubling time and ESR strength (Fig. 3*B*; Spearman, iESR ρ^2^ = 0.3144, *P* = 0.0066; Spearman, rESR ρ^2^ = 0.4942, *P* = 0.0003) as well as between degree of aneuploidy and ESR strength (Fig. 3*C*; Spearman, iESR ρ^2^ = 0.2864, *P* = 0.0103; Spearman, rESR ρ^2^ = 0.4707, *P* = 0.0004). It is worth noting that a few aneuploid strains were not able to mount the ESR, despite their slowed proliferation (i.e. strain A22 from Pavelka et al. (2010; 12); circled in red in Fig. 3). The strains that were unable to activate the ESR all harbored gains of chromosomes 2, 7, 11, 15, and 16. This observation suggests that some specific gene combinations prevent activation of the ESR despite slow proliferation. It will be interesting to determine the mechanism of this ESR suppression.

Finally, we probed for the existence of the CAGE signature in these complex aneuploid strains. The upregulated CAGE signature did not correlate with degree of aneuploidy (Supplemental Fig. 3*A*; Spearman, Upregulated CAGE ρ^2^ = 0.0679, *P* = 0.2416). We observed a correlation between the downregulated CAGE signature and degree of aneuploidy, but it was opposite to what would be expected if it were determined by degree of aneuploidy. Increased degree of aneuploidy correlated with increased expression of CAGE downregulated genes (Supplemental Fig. 3*A*; Spearman, Downregulated CAGE ρ^2^ = 0.2368, *P* = 0.0217). We conclude that the CAGE signature is not a common aneuploidy gene expression signature among complex aneuploid strains, but the ESR is.

### Proliferation rate determines ESR strength

The strong correlation between ESR strength and doubling times in complex aneuploid strains suggested that proliferation rate was the primary determinant of ESR strength. To directly test this possibility, we examined whether equalizing proliferation rate among complex aneuploid strains and euploid control strains affected the correlation between ESR strength and degree of aneuploidy by culturing cells in a phosphate-limited chemostat (4, 13). When proliferation rate was equalized in this manner, the ESR gene expression signature was no longer evident in aneuploid strains (Fig. 3*D*; Spearman, iESR ρ^2^ = 0.1912, *P* = 0.2066; Spearman, rESR ρ^2^ = 0.0107, *P* = 0.7850). We also probed for the existence of the CAGE signature in complex aneuploid strains grown under phosphate-limiting conditions. We observed no correlation between degree of aneuploidy and genes upregulated in the CAGE response and the opposite correlation as would have been expected for the downregulated genes of the CAGE signature (Supplemental Fig. 3*B*; Spearman, Upregulated CAGE ρ^2^ = 0.0030, *P* = 0.8916; Spearman, Downregulated CAGE ρ^2^ = 0.4364, *P* = 0.0.0438). We conclude that when proliferation is equally slow in euploid and aneuploid cells, the ESR caused by aneuploidy is no longer evident. This suggests that in both euploid and aneuploid cells, proliferation rate is the primary determinant of ESR strength.

### ESR induction in aneuploid cells causes ribosome loss

Tsai et al. (2019; 9) reported the CAGE gene expression signature as aneuploidy-specific and most similar to the hypo-osmotic stress response. This similarity is not surprising given that both the CAGE signature and the hypo-osmotic stress response are essentially oppositely regulated to the ESR. Given our findings that heterogeneous aneuploid populations do not exhibit a CAGE signature, we next determined whether aneuploid cells indeed experience hypo-osmotic stress that was proposed to occur in response to a decrease in cell density in aneuploid cells (9).

To assess induction of the hypo-osmolarity stress pathway, we probed activation of the hypo-osmolarity pathway MAP kinase Slt2 using a phospho-specific antibody that recognizes active phospho-Slt2 (14). Aneuploid cell populations showed a 2.22-fold increase in mean Slt2-phosphorylation compared to euploid control populations (Fig. 4*A* and *B*). The activation in the aneuploid cell populations was subtle compared to cell wall stress induced by prolonged Calcofluor White treatment (6.59-fold increase over euploid cell population). This difference in induction was not due to acute versus chronic induction of the hypo-osmolarity pathway because we treated cells with Calcofluor White for two hours before analyzing the phosphorylation state of Slt2. The subtle activation of the hypo-osmolarity pathway in aneuploid strains suggests the possibility that either all aneuploidies cause subtle activation of this stress pathway or that only a subset of aneuploid cells activates the pathway. We favor the latter possibility because we previously showed that not all aneuploidies cause cell wall defects (15).

**Figure 4.**
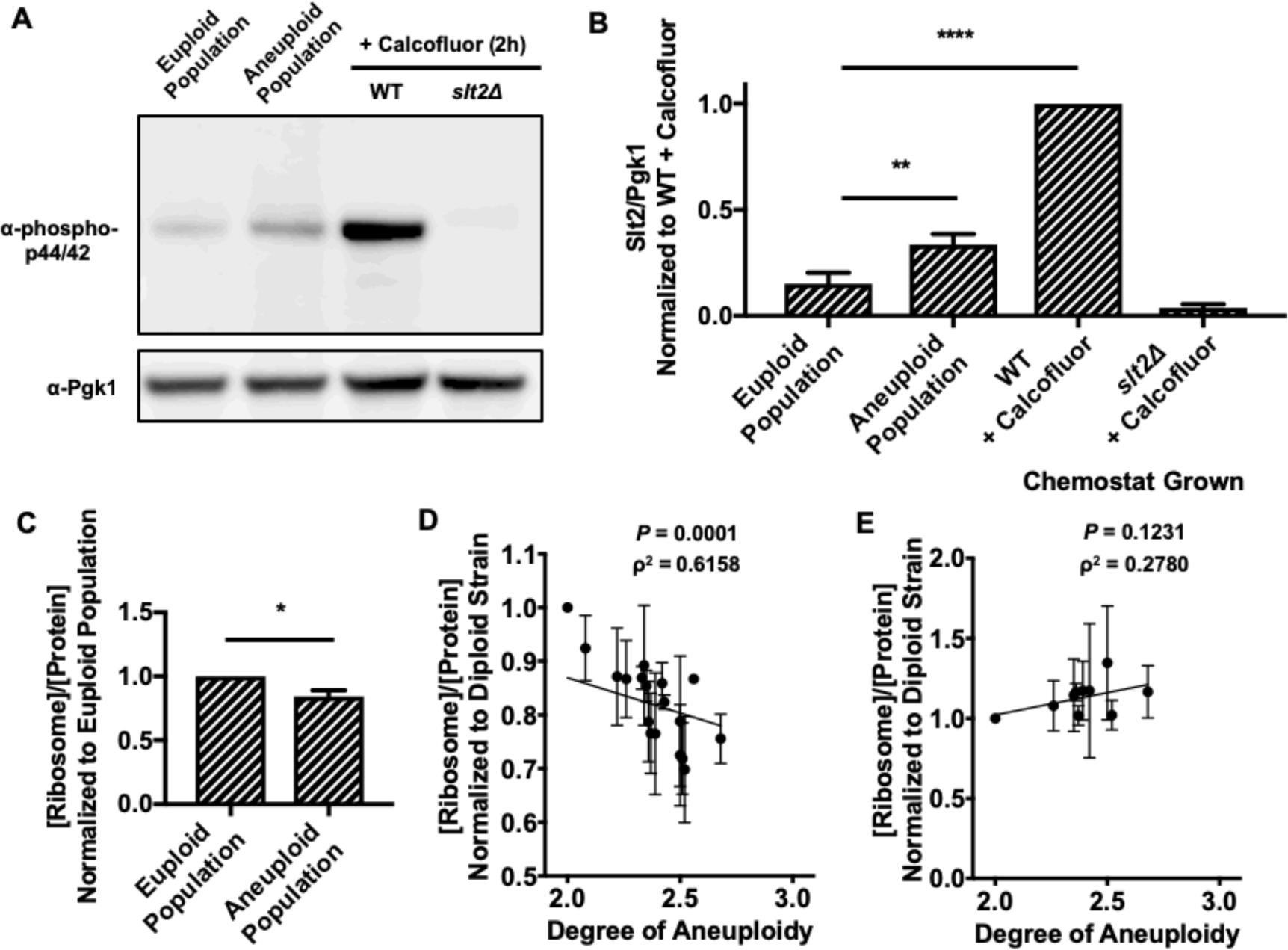
ESR induction causes ribosome loss in aneuploid strains. **(*A* and *B*)** Euploid and aneuploid cell populations were grown in YEPD with the 1:20 dilution protocol, and Slt2 Thr202/Tyr204 phosphorylation was determined. Wild-type euploid (A2050) and *slt2Δ* cells (A41265) treated with 5 μg/mL Calcofluor White for two hours served as positive and negative controls, respectively, in immunoblots (***A***). Pgk1 served as a loading control. Quantifications of Slt2 Thr202/Tyr204 phosphorylation are shown in (***B***). Slt2/Pgk1 values were normalized to the wild-type cells treated with Calcofluor White. Error bars represent standard deviation from the mean; one-way ANOVA test with multiple comparisons and Bonferroni correction, *P* < 0.0001 (****), *P* = 0.0021 (**). All other comparisons between samples had a significant difference of *P* < 0.0001 (****) with the exception of the euploid populations and *slt2Δ* + Calcofluor, which had a significant difference of *P* = 0.0288. **(*C*)** The fraction of ribosome in total protein extracts ([Ribosome]/[Protein]) was determined in euploid and aneuploid cell populations grown with the 1:20 dilution protocol. [Ribosome]/[Protein] in aneuploid cell populations was normalized to that in euploid cell populations. Error bars represent standard deviation from the mean; unpaired two-tailed t-test test, *P* = 0.0332 (*). **(*D*)** Aneuploid yeast strains harboring aneuploidies ranging from 2N to 3N were grown to log phase in YEPD and the fraction of ribosomes in total protein extracts ([Ribosome]/[Protein]) was determined. Correlation between [Ribosome]/[Protein] and degree of aneuploidy (ρ^2^ = 0.6158, *P* = 0.0001, Spearman) is shown. The calculated values were normalized to the [Ribosome]/[Protein] of a diploid control. **(*E*)** Aneuploid yeast strains harboring aneuploidies ranging from 2N to 3N were grown in a phosphate-limited chemostat and the fraction of ribosome in total protein extracts ([Ribosome]/[Protein]) was determined. Correlation between [Ribosome]/[Protein] and degree of aneuploidy (ρ^2^ = 0.2780, *P* = 0.1231, Spearman) is shown. The calculated values were normalized to the [Ribosome]/[Protein] of a diploid control. Error bars represent standard deviation from the mean.

The subtle activation of the hypo-osmolarity pathway in aneuploid cell populations was hard to reconcile with the comparatively dramatic effects on cell density reported to occur in aneuploid cell populations (9). Our observation that aneuploid cell populations exhibit the ESR provided an alternative hypothesis. A recent study by Delarue et al. (2018; 16) showed that the major determinant of cytoplasm density is the ribosomal fraction within a cell’s proteome. Given that aneuploid cell populations exhibit the ESR, which is characterized by the downregulation of ribosomal protein and ribosome biogenesis gene expression, we asked whether aneuploid cell populations harbor fewer ribosomes than euploid control populations. To address this question, we purified ribosomes from aneuploid and euploid populations grown using the 1:20 dilution protocol (*SI Appendix*, Fig. S1*B*).). This analysis revealed that ribosomes made up a significantly smaller fraction of total protein in aneuploid cell populations than in euploid cell populations (Fig. 4*C*).

Our analysis of complex aneuploid strains confirmed these results. Ribosome content inversely correlated with degree of aneuploidy (Fig. 4*D*; Spearman, ρ^2^ = 0.6158, *P* = 0.0001), which is consistent with ESR strength correlating with degree of aneuploidy. This correlation suggested that the ESR triggered ribosome loss and hence loss of cytoplasm density in aneuploid cells. If true, we would predict that equalizing proliferation rates among aneuploid and euploid cells should eliminate this correlation. This is what we observed. When we grew euploid and complex aneuploid strains in continuous culture under phosphate-limiting conditions, all strains not only exhibited similar ESR strengths, but ribosome content was no longer inversely correlated with degree of aneuploidy (Fig. 4*E*; Spearman, ρ^2^ = 0.2780, *P* = 0.1231). We propose that the decrease in cell density observed in aneuploid cells is caused by an ESR-induced loss of ribosomes.

## Discussion

Whether aneuploidy elicits a stereotypic transcriptional response in yeast and what this response may be has been controversial. The analysis of a series of disomic yeast strains harboring an additional copy of one of yeast’s 16 chromosomes showed that these strains exhibit the generic environmental stress response (ESR). However, other reports found this not to be the case. The analysis of five complex aneuploid yeast strains showed that only three out of these five strains exhibited the ESR (12). Most recently, Tsai et al. (2019; 9) reported that heterogeneous aneuploid cell populations also lack the ESR. We re-evaluated these reports to find that when a large number of complex aneuploid strains is analyzed and when gene expression profiles of mixed aneuploid cell populations are normalized to euploid populations that are in the same proliferation state as aneuploid populations, the ESR is evident. Together, these results indicate that aneuploid yeast strains exhibit the ESR. We consider this result not surprising given that the ESR is largely a consequence of slowed cell division (4, 8) and that aneuploidy causes proliferation defects (5). We further note that the ESR-like transcriptional signature is also observed in aneuploid mammalian cells (4). Together, these results indicate that the ESR and relatives of this signature are a pervasive response to aneuploidy.

In wild isolates of yeast that are naturally aneuploid, the ESR is less prevalent (1, 2). This observation indicates that under selective pressure, aneuploidies evolve and adaptation to aneuploidy-induced cellular stresses occurs such that proliferation rate is less affected by a particular chromosome gain or loss. Indeed, an aneuploidy tolerating natural gene variant has recently been described (2). Based on the analysis of two wild yeast isolates disomic for either chromosome 8 or 12, Hose et al. (2020; 2) proposed that the ESR in aneuploid cells is strongly influenced by the type of *SSD1* allele that a strain carries. Specifically, the ESR observed in aneuploid strains of the W303 background was attributed to the fact that W303 harbors a truncation allele of *SSD1*. The complex aneuploid strains and the aneuploid cell populations employed in our study are derivatives of the S288C strain background, which harbors a full-length *SSD1* allele. The finding that ESR strength correlates with degree of aneuploidy in these aneuploid S288C derivatives indicates that the ESR is not defined by *SSD1* allele identity but degree of aneuploidy. It is nevertheless possible that mutations and specific aneuploidy combinations exist that suppress the ESR gene signature. Indeed, we found that strains harboring a gain of chromosomes 2, 7, 11, 15, and 16 do not exhibit the ESR despite proliferating slowly. Understanding why these chromosome combinations suppress the ESR will be interesting. We speculate that growth-promoting pathways known to negatively regulate the ESR, such as the PKA and TOR pathways, are hyperactive in these strains.

Our data indicate that the previously described aneuploidy-specific CAGE gene expression signature is an artifact caused by normalizing the gene expression of actively dividing aneuploid cells to that of euploid control cells that had grown to stationary phase. We show that the ESR in the euploid control populations that had grown into stationary phase was stronger than the ESR observed in aneuploid cell populations that, due to their poor proliferation, were still actively proliferating. Thus, when euploid stationary phase cells were used for normalization, aneuploid cells exhibited the CAGE signature in which many ESR genes are oppositely regulated. As such, it is not surprising that the CAGE signature is most similar to the previously described hypo-osmolarity gene expression signature. Gasch et al. (2000; 6) identified two stresses that do not result in activation of the ESR: cold shock and hypo-osmotic shock. Under these stresses, the ESR is oppositely regulated. In particular the rESR, which encompasses genes encoding transcription and translation factors and ribosomal proteins are upregulated rather than downregulated (6). While it is not clear why ribosome production must be upregulated during cold shock, we understand why this occurs during hypo-osmotic shock. During hypo-osmotic shock, water uptake increases (10). To avoid cytoplasm dilution during this water influx, production of ribosomes, which are the main determinants of cytoplasm density (16), must be upregulated or at least prevented from being downregulated. Further analysis of these exceptional stress conditions, during which the rESR is not downregulated will provide key insights into regulation of this central stress and slow proliferation response.

Under most, if not all stress conditions, cell proliferation is slowed or halted, and the ESR signature is evident with the exceptions noted above. Downregulation of the rESR generally correlates much better with degree of slow proliferation than induction of the iESR. This is not surprising. The vast majority of the genes that are part of the rESR are involved in transcription and translation. Cell size growth, or cellular enlargement, needs to be attenuated during any stress that causes a slowing or halting of cell proliferation to prevent cells from growing too large. Were this to occur, DNA becomes limiting, causing numerous defects including impaired cell proliferation, cell signaling, and gene expression (17, 18). In contrast, the iESR is aimed at alleviating cellular stress which requires expression of genes unique to specific stresses rather than control of biomass production.

Our results show that repression of rESR genes in response to aneuploidy has profound consequences on cellular proteome composition. It leads to a significant drop in the contribution of ribosomes to the cell’s total protein. This not only leads to downregulation of translational capacity but, because ribosomes are the key determinant of cytoplasmic density (16), is likely the major cause of loss in cellular density previously reported to occur in aneuploid cells (9). It thus appears that activation of the ESR in aneuploid cells serves two purposes. It protects cells from cellular stresses caused by an unbalanced genome and prevents excessive cellular enlargement during their slowed cell cycles. Understanding how slowed proliferation leads to activation of the ESR will provide critical insights into the coordination between cell division and macromolecule biosynthesis.

## Acknowledgements

Thank you to Summer Morrill, Xiaoxue Zhao, and David Waterman for comments and the MIT BioMicroCenter for RNA-Seq. This work was supported by NIH grant R35 GM118066 to A.A., who is an investigator of the Howard Hughes Medical Institute, the Paul F. Glenn Center for Biology of Aging Research at MIT and the Ludwig Center at MIT. AK was supported in part by NHGRI grant T32 HG-00035. The research of MJD was supported in part by a Faculty Scholar grant from the Howard Hughes Medical Institute and by NIGMS grant P41 GM103533. MJD is a Senior Fellow in the Genetic Networks program at the Canadian Institute for Advanced Research.

## Materials and Methods

### Dataset Processing and Single-Sample Gene Set Enrichment Analysis

Raw RNA-Seq data was obtained as described below or through download from Tsai et al. (2019; 9) with gene accession number GSE107997. Reads were aligned to a *S. cerevisiae* transcriptome with STAR version 2.5.3a (19) and gene expression was quantified with RSEM version 1.3.0 (20). Transcript per million (TPM) values were offset by +1, log_2_ transformed, and used to prepare GCT files for single-sample Gene Set Enrichment Analysis (ssGSEA, version 7.7; 21, 22) obtained from the Indian University public GenePattern server (23). ssGSEA projections were prepared for the Environmental Stress Response (ESR) originally described by Gasch et al. (2000; 6) “common aneuploidy gene-expression” (CAGE) signatures identified in Tsai et al. (2019; 9). Sort order has a subtle effect on ssGSEA.

### Differential Gene Expression Analysis

Raw RNA-Seq data was obtained through download from Tsai et al. (2019; 9) with gene accession number GSE107997. Integer count values derived from RSEM processing were used as input to differential expression analysis with DESeq2 (version 1.24.0; 24) using normal log fold change shrinkage. Expression data from aneuploid cell populations generated by tetrad dissection were pooled to create “aneuploid populations (Tetrad)”. Expression data from aneuploid populations obtained from *MATa* selection were pooled to create “aneuploid populations (*MATa* Selection)”. These populations, as well as both the euploid populations obtained from tetrad dissection and *MATa* selection, were compared to the exponentially growing haploid control. Differential expression data was visualized using TIBCO Spotfire Analyst version 7.11.1.

### Stationary-Phase Growth Timecourse

S288C wild-type cells (A2050) were inoculated into 25 mL YEPD and grown overnight at 25 °C. After approximately 12-14 hours of growth, cells were diluted to OD(600nm) = 0.1 with YEPD + 2% glucose and grown for an additional 4 hours at 25 °C. Once the (OD600nm) reached approximately OD(600nm) = 0.3, samples for transcriptomics were taken and OD(600nm) was measured every 2 hours for 28 hours. Optical Density was measured at 600 nm (OD(600nm)) with a spectrophotometer.

### Heterogeneous Euploid and Aneuploid Population Generation Using the Tsai et al. (2019; 9) Protocol

*pRS315-STE2pr-spHIS5* S288C diploids (A40877) and triploids (A40878) (*SI Appendix*, Table S1) were grown overnight in SD - LEU medium and subsequently sporulated in “Super Sporulation Medium” (1% potassium acetate and 0.02% raffinose) from Tsai et al. (2019). Sporulated tetrads were then dissected. (Note in the publication by Tsai et al. (2019), sporulated tetrads were also *MATa* selected through histidine prototrophy. We did not generate cell populations in this manner). Individual colonies were grown for 14-16 hours in 200 μL of YEPD + 2% glucose in a 96 deep-well plate. 300 μL of YEPD were then added to cultures. The cultures were grown for 5 additional hours, pooled, and samples for RNA-Seq and [Ribosome]/[Protein] content measurements were taken.

### Heterogeneous Euploid and Aneuploid Population Generation Using the 1:20 Dilution Protocol

*pRS315-STE2pr-spHIS5* S288C diploids (A40877) and triploids (A40878) (*SI Appendix*, Table S1) were grown overnight in SD - LEU medium and subsequently sporulated in the “Super Sporulation Media” mentioned above. Sporulated tetrads were then tetrad dissected. Individual colonies obtained from spores were grown for 14-16 hours in 200 μL of YEPD in a 96 deep-well plate. Cultures were then diluted 1:20 in YEPD + 2% glucose, grown for another 5 hours, then pooled, and diluted to approximately OD(600nm) = 0.3. The pooled aneuploid populations were grown for an additional 2 hours, and samples for RNA-Seq, [Ribosome]/[Protein] content measurements, and Slt2 phosphorylation state analysis were taken.

### Growth Conditions for Complex Aneuploid Strains

Complex aneuploid strains were generated by Pavelka et al. (2010; 12) (*SI Appendix*, Table S2). Complex aneuploid strains were grown on YEPD + 2% glucose plates for 2 days at 25 °C. Colonies were inoculated overnight in 25 mL YEPD + 2% glucose and grown at 25 °C. After 12-14 hours of growth at 25 °C, cells were diluted to approximately OD(600nm) = 0.1 and grown for an additional 4 hours at 25 °C. Cells were then harvested for RNA-Seq and [Ribosome]/[Protein] content measurements.

### Doubling Time Measurements

Doubling time of complex aneuploid strains were measured in a 96 well format in YEPD + 2% glucose at 25 °C. OD(600nm) values were taken in 20 minute intervals over 5 hours, and doubling time was calculated from the growth curves generated.

### Growth in Phosphate-Limiting Chemostats

Selected complex aneuploid strains were grown on YEPD + 2% glucose plate for 2 days at 30 °C. Strains were inoculated in phosphate-limited media (25) and grown overnight. Once chemostats were set up and filled, phosphate-limited media in the chemostat was inoculated with 2 mL of overnight culture, and cells were allowed to grow for 30-36 hours although some strains required 48+ hours of growth. The dilution pumps were then turned on at a dilution rate of 0.11 +/- 0.01 chemostat volumes per hour. The chemostat was sampled daily to measure effluent volume, hemocytometer counts, and OD(600nm) measurements. When growth in the chemostat had reached a steady-state, defined by less than 5% change from the previous day’s measurements, samples were harvested for RNA-Seq and [Ribosome]/[Protein] content.

### RNA-Seq

3-5 mL samples of culture were taken, spun down at 3000rpm for 5 minutes, washed with 1 mL DEPC water, and transferred to a 2 mL RNase-free screw-cap tube. Samples were spun again at 8000rpm for 3 minutes, and supernatant was aspirated. Cells were snap frozen with liquid nitrogen and stored at -80 °C. RNA samples were prepared with RNeasy mini kit from Qiagen and treated with DNase on-column treatment (RNase-free) from Qiagen. Purified RNA was used in two different library preparation methods. In experiments with complex aneuploid strains, total RNA was sequenced using Illumina Truseq followed by Nextera or Roche KAPA. In all other experiments, total RNA was sequenced using Illumina Truseq followed by Roche KAPA. All sequencing was done using an Illumina HiSeq2000.

### [Ribosome]/[Protein] Content Measurements

50 mL samples of culture were taken, spun down at 3000rpm for 5 minutes, frozen with liquid nitrogen, and stored at -80 °C. Cells were resuspended in 30 mL lysis buffer (20mM HEPES pH 7.4, 100mM potassium acetate, 2mM magnesium acetate, 3mM DTT, and EDTA-free protease inhibitor (Roche, 11836170001)), and 0.5 mg/mL zymolyase was added. Resuspended cells were lysed twice with a French Press. Lysed samples were then spun at 19,000rpm for 20 minutes at 4 °C to remove cell debris. Protein concentration of the cell lysate ([Protein]) was measured in triplicate by Bradford Assay.

10 mL cell lysate was overlaid onto 15 mL pre-chilled (4 °C) sucrose solution (30% sucrose, 20mM HEPES pH 7.4, 500mM potassium acetate, 2mM magnesium acetate, and 3mM DTT), which was then spun at 50,000 rpm for 4 hours at 4 °C. Solution above assembled ribosome pellet was poured from the tube after spin, and the tubes were dried upside-down for 10 minutes to remove excess liquid. Pellets were resuspended with 1 mL lysis buffer, and absorbance at 260nm was measured with a Nanodrop to obtain concentration of purified assembled ribosomes ([Ribosome]).

### Calcofluor Treatment

S288C wild-type (A2050) and *slt2Δ* (A41265) cells were inoculated into 25 mL YEPD + 2% glucose, and grown at 25 °C for 12-14 hours. Cells were diluted to approximately OD(600nm) = 0.1, grown for an additional 4 hours, and treated with 5 μg/mL Calcofluor White for 2 hours. Samples were harvested for Western Blot analysis.

### Western Blot and Quantification

1 OD(600nm) unit of sample was Trichloroacetic acid (TCA) precipitated. 20 μL of each sample was run on a NuPAGE 4-12% Bis-This protein gel from Invitrogen and then transferred to PVDF membrane from EMD Millipore. Phospho-p44/42 MAPK (Thr202/Tyr204) antibody (1:1000; Cell Signaling Technology, #9101) was used to detect phosphorylated Slt2. Pgk1 (Pgk1 antibody, 1:4000; ThermoFisher, 22C5D8) was used as a loading control. Immunoblots were incubatedwith HRP-conjugated secondary antibodies and ECL Western Blotting Detection Reagents from Amersham and then scanned on an ImageQuant LAS4000. Signal was quantified on an ImageQuant LAS4000 and integrated densities of bands quantified using ImageJ. Three separate immunoblots were quantified and normalized to wild-type cells treated with Calcofluor.

---

**Figure S1.**
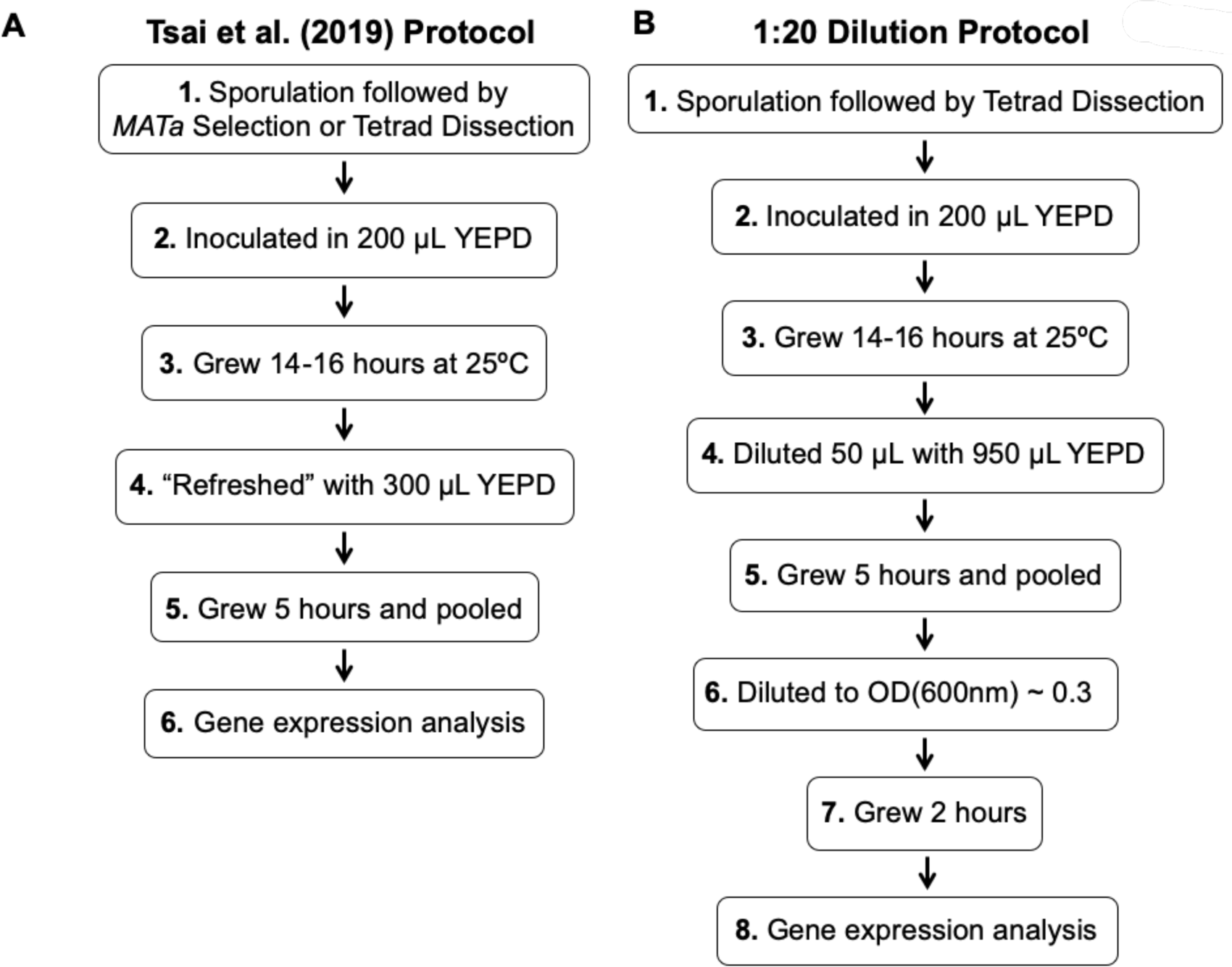
Generation of heterogeneous aneuploid populations. **(*A*)** Protocol developed by Tsai et al. (2019; 9) to generate aneuploid cell populations. Cells harboring random aneuploidies were generated by sporulation of *pRS315-STE2pr-spHIS5* S288C triploids (A40878) and subsequent tetrad dissection or *MATa* selection through histidine prototrophy. Individual colonies were grown for 14-16 hours in 200 μL of YEPD in a 96 deep-well plate. 300 μL of YEPD were then added to cultures. The cultures were grown for 5 additional hours, pooled, and analyzed. **(*B*)** Protocol to avoid growth of cell populations into stationary phase (1:20 dilution protocol). Cells harboring random aneuploidies were generated by sporulation of *pRS315-STE2pr-spHIS5* S288C triploids (A40878) and subsequent tetrad dissection. Individual colonies were grown for 14-16 hours in 200 μL of YEPD in a 96 deep-well plate. Cultures were then diluted 1:20 in YEPD, grown for another 5 hours, then pooled, and diluted to approximately OD(600nm) = 0.3. The pooled aneuploid populations were grown for an additional 2 hours, and samples were taken.

**Figure S2.**
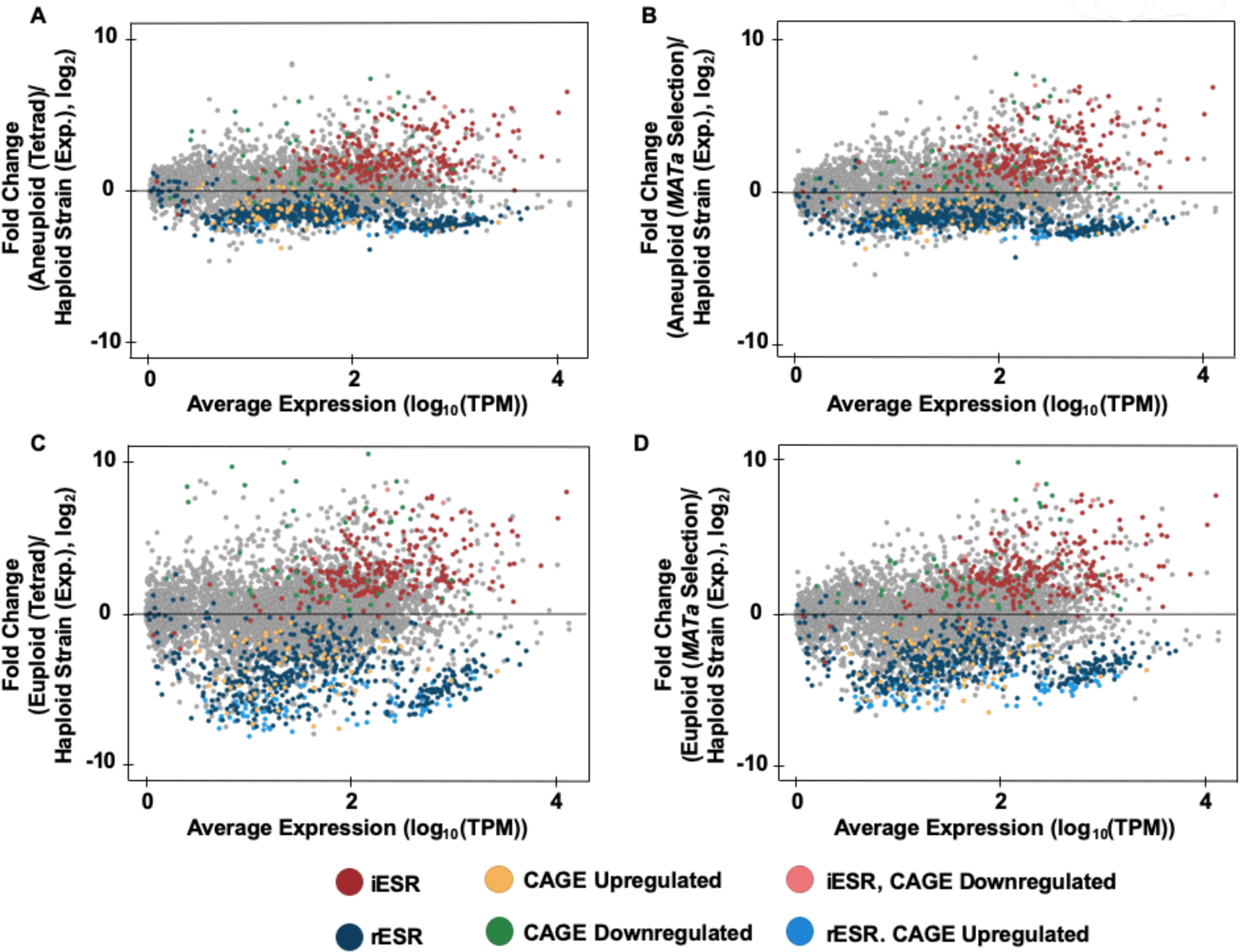
Comparison of aneuploid and euploid gene expression patterns from Tsai et al. (2019; 9) to an to exponentially growing haploid strain. RNA-Seq data from Tsai et al. (2019; 9) were processed using the Expectation Maximization (RSEM) method. Transcript per million (TPM) values were calculated and log_2_ transformed. Expression data from aneuploid cell populations generated by tetrad dissection were pooled to create “aneuploid populations (Tetrad)”. Expression data from aneuploid populations obtained from *MATa* selection were pooled to create “aneuploid populations (*MATa* Selection)”. The x axis shows log_10_(average basal expression), and the y axis shows differential expression between euploid or aneuploid populations (Tetrad and *MATa* Selection) and an exponentially growing haploid strain (Tsai et al. (2019; 9), accession number: GSE107997). Colors specified refer to iESR, rESR, CAGE upregulated, CAGE downregulated, and those iESR genes downregulated in the CAGE signature and rESR genes upregulated in the CAGE signature. Differential expression graphs are shown for aneuploid cell populations (Tetrad) compared to the exponentially growing haploid strain **(*A*)**, aneuploid cell populations (*MATa* Selection) compared to the exponentially growing haploid strain **(*B*)**, euploid cell population (Tetrad) compared to the exponentially growing haploid strain **(*C*)**, and euploid cell population (*MATa* Selection) compared to the exponentially growing haploid strain (RLY4388 **) (*D*)**.

**Figure S3.**
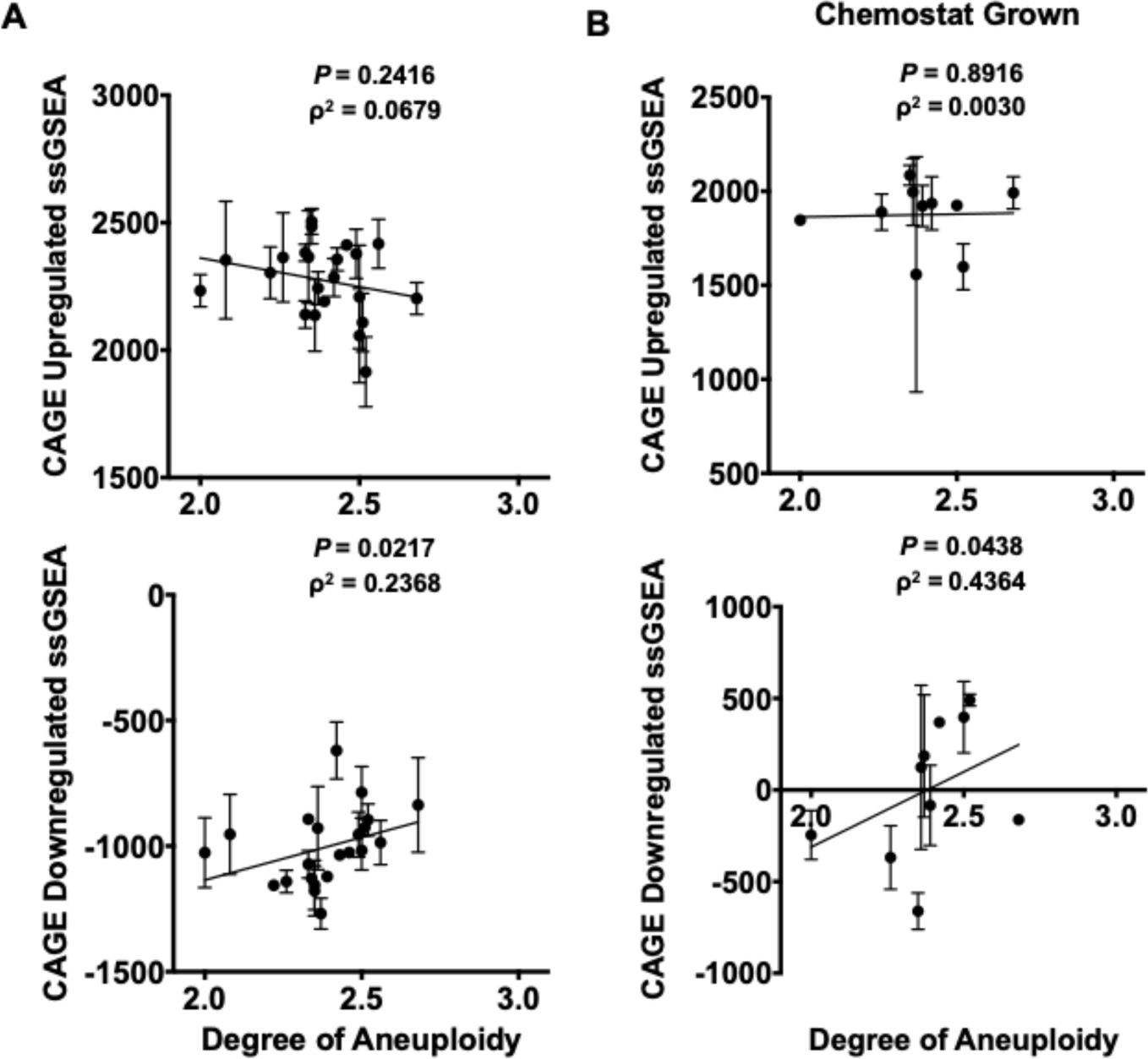
Correlation between growth rate and the CAGE gene expression signature in complex aneuploid strains. Transcriptomes of the complex aneuploid strains were analyzed by RNA-Seq, and ssGSEA projection values were calculated for CAGE upregulated and CAGE downregulated genes. **(*A*)** Correlation between CAGE upregulated ssGSEA projections and degree of aneuploidy (Spearman, ρ^2^ = 0.0679, *P* = 0.2416) and CAGE downregulated ssGSEA projections and degree of aneuploidy (Spearman, ρ^2^ = 0.2368, *P* = 0.0217) in complex aneuploid strains grown in YEPD. **(*B*)** Correlations between CAGE upregulated ssGSEA projections and degree of aneuploidy (Spearman, ρ^2^ = 0.0030, *P* = 0.8916) and CAGE downregulated ssGSEA projections and degree of aneuploidy (Spearman, ρ^2^ = 0.4364, *P* = 0.0438) grown in a phosphate-limited chemostat. Error bars represent standard deviation from the mean.

**Table S1.** Euploid strains. Description of the strain names, genotypes, and source used in this paper.

| Strain Name | Genotype | Source |
| --- | --- | --- |
| A2587 | W303, MATa ade2-1, leu2-3, ura3, trp1-1, his3-11,15, can1-100, GAL, psi+ | Kim Nasmyth |
| A2050 | S288C, MATa ura3-52 his3Δ leu2-3 112 trp1Δ | Frank Luca |
| A41265 | BY4741, MATa his3Δ leu2Δ met15Δ ura3Δ slt2Δ::KanMX | BY4741 Deletion Collection |
| A40877 | BY4743, MATa/$\alpha$ his3Δ/his3Δ leu2Δ/leu2Δ met15Δ/MET15 ura3Δ/ura3Δ lys2Δ/LYS2 pRS315-STE2pr-spHIS5 | Tsai et al. (2019; 9) RLY9593 |
| A40878 | WT Triploid, MATa/a/$\alpha$ his3Δ/his3Δ/his3Δ leu2Δ/leu2Δ/leu2Δ met15Δ/met15Δ/MET15 ura3Δ/ura3Δ/ura3Δ lys2Δ/LYS2/LYS2 pRS315-STE2pr-spHIS5 | Tsai et al. (2019; 9) RLY9596 |

**Table S2.**
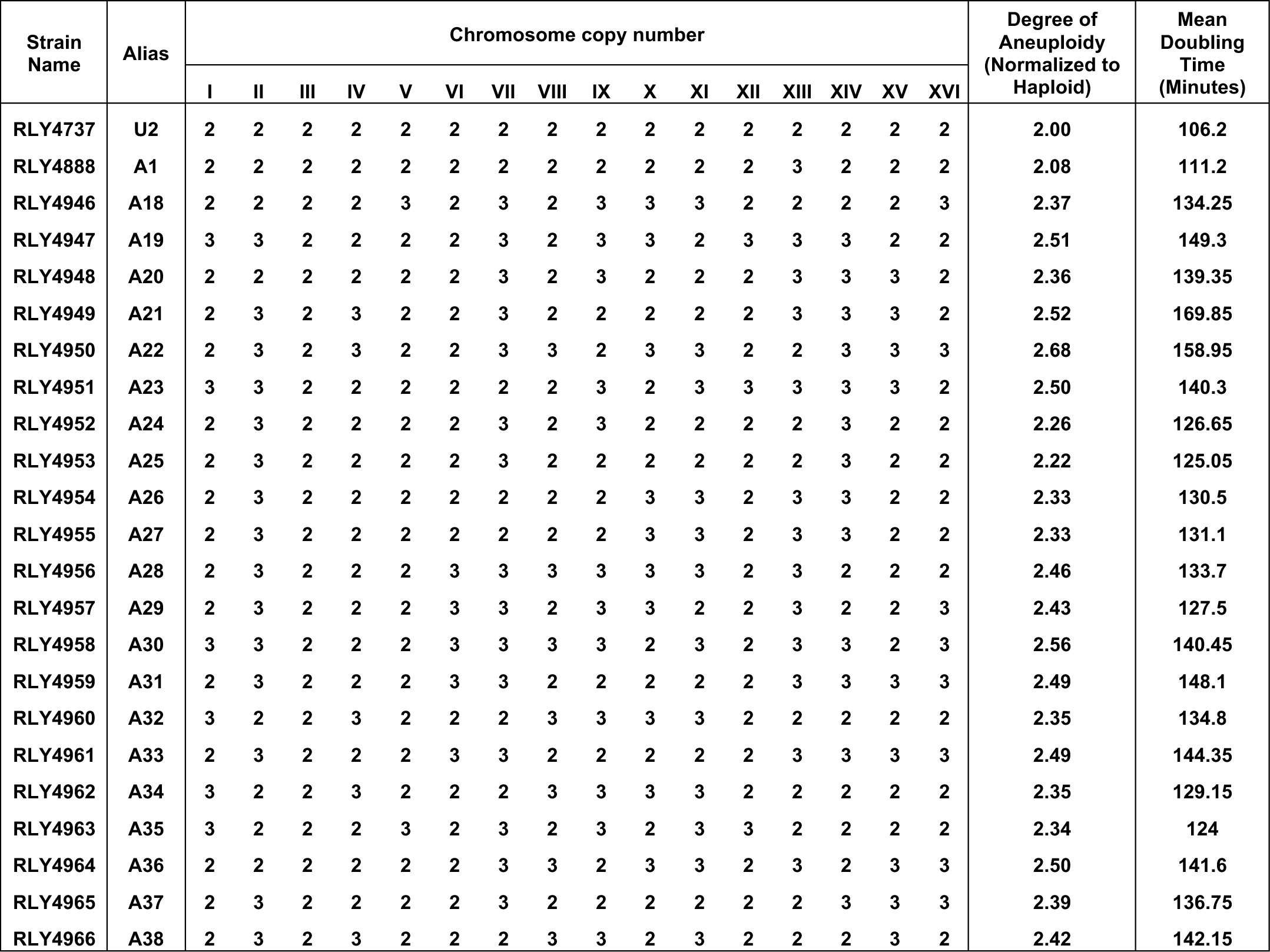
Complex aneuploid strains. Description of the strain names, aliases, karyotypes and mean doubling times of complex aneuploid strains generated by Pavelka et al. (2010; 12).

